# De novo clustering of large long-read transcriptome datasets with isONclust3

**DOI:** 10.1101/2024.10.29.620862

**Authors:** Alexander J. Petri, Kristoffer Sahlin

## Abstract

Long-read sequencing techniques can sequence transcripts from end to end, greatly improving our ability to study the transcription process. Although there are several well-established tools for long-read transcriptome analysis, most are reference-based. This limits the analysis of organisms without high-quality reference genomes and samples or genes with high variability (e.g., cancer samples or some gene families). In such settings, analysis using a reference-free method is favorable. The computational problem of clustering long reads by region of common origin is well-established for reference-free transcriptome analysis pipelines. Such clustering enables large datasets to be split up roughly by gene family and, therefore, an independent analysis of each cluster. There exist tools for this. However, none of those tools can efficiently process the large amount of reads that are now generated by long-read sequencing technologies.

We present isONclust3, an improved algorithm over isONclust and isONclust2, to cluster massive long-read transcriptome datasets into gene families. Like isONclust, IsONclust3 represents each cluster with a set of minimizers. However, unlike other approaches, isONclust3 dynamically updates the cluster representation during clustering by adding high-confidence minimizers from new reads assigned to the cluster and employs an iterative cluster-merging step. We show that isONclust3 yields results with higher or comparable quality to state-of-the-art algorithms but is 10-100 times faster on large datasets. Also, using a 256Gb computing node, isONclust3 was the only tool that could cluster 37 million PacBio reads, which is a typical throughput of the recent PacBio Revio sequencing machine.

**Availability:** https://github.com/aljpetri/isONclust3

## 1 Introduction

Long-read transcript sequencing using PacBio and Oxford Nanopore Technologies (ONT) has been key in investigating complex isoform landscapes across various organisms. There exist several genome and annotation-based tools capable of analyzing transcriptomic long-read sequencing data Orabi *et al*. (2023); Prjibelski *et al*. (2023); Kovaka *et al*. (2019); Tung *et al*. (2019), but those tools are limited to well-sequenced and annotated organisms, and can, at best, be used with references from closely related organisms. Additionally, such tools do not work well for data that does not match the reference, e.g., gene families not well represented in current reference genomes Sahlin *et al*. (2018) or cancer samples with high diversity to the reference. This has created a need for de novo transcriptome analysis. The generally established workflow of de novo transcriptome analysis is to cluster the reads into transcripts, genes, or gene families, then perform read error correction (if needed), and finally reconstruct transcripts present in the individual clusters de la Rubia *et al*. (2022); Sahlin and Medvedev (2020, 2021); Petri and Sahlin (2023). The most resource-intensive step of the three is clustering, as the two downstream steps can work with sub-instances of the data (the clusters). However, the data yields for long-read sequencing techniques have been greatly enhanced over the last couple of years Marx (2023). The order of magnitude increase in data poses scaling challenges to existing state-of-the-art algorithms. While several different algorithms exist for the de novo clustering of generic protein Li and Godzik (2006); Paccanaro *et al*. (2006); Steinegger and Söding (2017), 2018) and nucleotide sequences Li and Godzik (2006); Edgar (2010); James *et al*. (2018); Ghodsi *et al*. (2011), transcripts have distinguished characteristics such as isoforms stemming from the same gene but having different splice patterns that set this problem apart from the other sequence clustering problems. Additionally, long-read transcriptome sequencing data pose challenges such as 3’ and 5’ transcript and sequencing variability, RNA degradation, and variable error rates between and within the reads.

### 1.1 Previous work on clustering of long-read transcriptome data

Several *de novo*-based clustering methods have been proposed (ToFU Gordon *et al*. (2015), IsoCon Sahlin *et al*. (2018), CARNAC-LR Marchet *et al*. (2018), isONclust Sahlin and Medvedev (2020), RATTLE de la Rubia *et al*. (2022), GeLuster Ma *et al*. (2024)). However, early tools such as ToFU and IsoCon Sahlin *et al*. (2018) were initial tools designed only for early PacBio datasets containing in the order of thousands to tens of thousands of reads. CARNAC-LR, RATTLE, GeLuster, isONclust were introduced and shown to handle hundreds of thousands up to a couple of million of reads. The CARNAC-LR algorithm creates a similarity graph using an all-vs-all read mapper such as minimap2 Li (2018) and derives an upper bound for a number of clusters in the graph. In the next step boundaries for initial clusters are refined to fulfill a partitioning condition to yield the final clustering. In CARNAC-LR, all-vs-all mapping is a bottleneck and the support for CARNAC-LR has been discontinued Marchet (????). We have previously introduced isONclust Sahlin and Medvedev (2020), which is based on minimizers. IsONclust is a greedy algorithm that sorts the reads in decreasing order by a combination of their quality and length. In the second step, reads are iterated over in the sorted order and the first read forms a cluster and subsequently all reads are compared to existing clusters, using minimizers Roberts *et al*. (2004). If a read has high similarity with an existing cluster, the read is added to the cluster, if not, the read forms a new cluster. Each cluster is represented by the minimizers of the read that initialized the cluster. IsONclust has widely been used over recent years both for transcriptome Kumazawa *et al*. (2022); Walter and Puniamoorthy (2022); Westrin *et al*. (2024); Unneberg *et al*. (2024) and amplicon sequencing analysis Sahlin *et al*. (2021); Pomerantz *et al*. (2022). However, it is not able to cluster reads transcribed in different orientations, and the algorithm is, by today’s standards, slow and memory intense. A more integral issue with the algorithm is that isONclust uses only the minimizers from the initial read forming a new cluster as the cluster’s representation. Therefore, the cluster representation is static and this prevents reads from isoforms with larger difference in exon structures to be placed in the same cluster. ONT, in collaboration with us, reimplemented the isONclust algorithm in the C++ programming language to improve its runtime and added some optional features naming it isONclust2 isONclust2 (2020). IsONclust2 employs the same algorithm as isONclust, but has in independent evaluation demonstrated about 3.5-4 times speedup over isONclust Sagniez *et al*. (2024), although at the cost of a high memory usage, as we show in our benchmark. IsONclust2 was, however, implemented such that it requires to be run within a snakemake wokflow, hindering usability of the tool. In addition, the tool has been discontinued and is no longer supported isONclust2 (2020). The RATTLE algorithm works similarly to isONclust in a greedy fashion, but with the differences that reads are sorted only by length in decreasing order in the first step. In the clustering step, RATTLE uses all *k*-mers to compare reads to existing clusters. Both isONclust and RATTLE have scalability issues, as we demonstrate. GeLuster is a recently published algorithm. In a first step, it selects a subset of the reads with a minimum length requirement (called pseudo-references), sharing low identity with each other by iterating through the reads in sorted order. In a second step, it aligns the remaining reads to the pseudo-references to form pre-clusters. The reads that have not been aligned properly to any of the pseudo-references will be passed as the input to the next iteration where the length requirement for being a pseudo-reference is reduced. This algorithm is performed iteratively with 3 iterations set as default. It has shown to improve the clustering while requiring less computational resources than isONclust, and RATTLE Ma *et al*. (2024). However, none of these algorithms are able to scale to datasets with tens to hundreds of millions of reads, as is common for the latest PacBio Revio or ONT PromethION devices. For example, isONclust is a core component in de novo transcriptomics pipelines used by companies such as biobam biobam (2022). Biobam was unable to process such large datasets with any of the algorithms (personal communication with Biobam), which we also demonstrate in this study using a PacBio Revio dataset. This prompted further development of algorithms to cluster currently generated long-read transcriptome sequencing datasets into gene families.

## 1.2 Our contribution

We developed isONclust3, an algorithm designed to address the shortcomings in scalability and accuracy of isONclust and isONclust2. Similarly to isONclust (and isONclust2), isONclust3 initially employs a greedy clustering algorithm using a minimizer-based approach to estimate similarity and to represent clusters. However, isONclust3 addresses the accuracy, time, and memory limitations with isONclust and other algorithms mentioned above in the following ways. Unlike isONclust and other approaches, isONclust3 updates the cluster representations during clustering by adding *high-confidence minimizers* from new reads assigned to the cluster. High-confidence minimizers are inferred from the quality values of the base calls in the minimizer. Such cluster updating is needed to accommodate new information, from e.g., additional exons, SNP variation from alleles, intron retentions, or other transcriptomic variability belonging to the gene families. The dynamic updating with high-confidence minimizers enables isONclust3 to cluster more reads from the same gene family, while keeping the number of minimizers stored for each cluster informative in comparison to adding all minimizers of a read (as for isONclust 1 and 2) when clustering reads from error prone sequencing technologies. Also, in addition to the greedy clustering in isONclust and isONclust2, isONclust3 can employ an efficient iterative cluster-merging step where clusters can be merged if they share enough high-confidence seeds. This step is optional to isONclust3 and it can improve clustering quality significantly over the initial clustering step on noisier ONT reads. Additionally, since genes can express transcripts in both directions on the genome, isONclust3 implements both non-canonical and canonical minimizers as seeds (unlike isONclust), to tailor for the scenario to keep these in separate and the same cluster, respectively. We tested isONclust3 and other state-of-the-art tools on ONT and PacBio data from different organisms as well as on simulated data. IsONclust3 shows slightly better or comparable clustering quality to other tools and is in many cases 1-2 orders of magnitude faster than other tools with comparable or smaller memory footprint. Most importantly, we demonstrate that isONclust3 is the only tool that can cluster data from the PacBio Revio sequencing machine using a 256Gb computing node.

## 2 Methods

### 2.1 Preliminaries

#### 2.1.1 Canonical minimizers

Let *r ∈ R* denote a string consisting of letters in Σ= *{A, C, G, T}*, that represents a sequencing *read*. We use zero-indexed notation for strings and let *r*[*i, i* + *k*) denote a substring (*k-mer*) on *r*, covering positions from *i* to *i*+*k −*1. Let a hash function *h* induce a *total ordering* over the *k*-mers. Given two integers *k* and *w* with 1 *≤ k ≤* |*r*| and 1 *≤ w ≤* |*r*| *− k* + 1, the (*k, w*)-minimizer Roberts *et al*. (2004) *m* in an interval *r*[*i, i* + *w*), referred to as a *window*, is the smallest *k*-mer starting in the window. Repeating *k*-mers within a window are resolved by choosing the leftmost *k*-mer. We refer to a (*k, w*)-minimizer as a minimizer when *k* and *w* are not important for context.

A canonical minimizer is a minimizer formed from taking the smallest *k*-mer among both the forward and the reverse complement *k*-mers in a substring *s ≥ k* of *r*, where *s* = *r*[*i, i* + *w* + *k −* 1]. We will refer to minimizers and canonical minimizers as seeds, as our approach work with any *k*-mer construct used for similarity search such as syncmers Edgar (2021), or when using all *k*-mers.

#### 2.1.2 High-confidence seeds

Base call qualities are measured for each base by the Phred quality score Ewing *et al*. (1998) (Q score) that can be transformed to a probability *q*_*i*_ that the base is correctly called through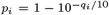, where *p*_*i*_ ∈ [0, 1] of the base located at position *i* in *r*. We compute the confidence of a seed spanning positions *r*[*i, i*+*k*) as 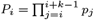. The formula computes the probability (as estimated by quality values) of the *k*-mer being correctly base-called in all of its nucleotides. If *Q*_*i*_ is greater than a threshold *T*_1_, we denote the seed as a *high-confidence seed* (HCS). We set *T*_1_ depending on the sequencing technology (analysis in section 3.1).

### 2.2 Algorithm

We describe the three steps of the algorithm below. Pseudocode of the clustering step is found in Suppl. Section S1 and a time complexity analysis in Suppl. Section S2.

#### Sorting

For each read we generate canonical minimizers and their associated qualities *Q*. We then sort the reads in decreasing order with respect to the number of HCS. This is in contrast to isONclust, which sorts according to the expected number of error-free *k*-mers in a read *r* and is not directly correlated to the amount of informative seeds used in the clustering. IsONclust3’s read ordering is therefore directly linked to the amount of useful information a read carries for the clustering. The sorted reads are stored in a file on disk.

#### Clustering

In the second step, the sorted reads are streamed over. The first read initializes a cluster and the HCS of the read are stored as a representation of the cluster. Consecutive reads are then compared against existing clusters in the following manner. If a read does not share a sufficient fraction (threshold *T*_2_) of (all) seeds from the read with the HCS of any existing cluster, it forms a new cluster. Only the HCS of the read kept as the cluster representation. Otherwise, if a read shares enough seeds with the HCS of at least one cluster, it is added to the cluster with the most shared seeds, and any HCS of the read not already existing in the cluster are added to the HCS of the cluster. For the clustering, isONclust3 stores a hash table that associates seeds as keys with arrays of cluster identifiers. For a visualization of the clustering approach, see Suppl. Fig. 1. The pseudocode of the algorithm is presented in Algorithm **??**. The on-the-fly updated cluster representations accommodate HCS stemming from additional exons, SNV variations, or other transcriptomic variability. This is in contrast to isONclust, which only uses the minimizers of the initial read as the cluster representation.

#### Merging clusters via cluster merging

isONclust3 also has the option to iteratively merge clusters after the greedy clustering step that can be activated via `–post-cluster`. Once clusters have been generated from sorted reads as described above, the clusters themselves have accumulated HCS along the way. A look-up table with cluster ID as key and a vector of HCS is constructed. The iterative cluster merging then visits clusters in ascending order of size (number of HCS) and finds other clusters with the highest overlap of HCS. This is done in a similar manner to the initial clustering, only that now clusters are compared, and the size ordering matters. Specifically, clusters can only be merged into larger clusters. If a (smaller) cluster *c*_1_ has the highest number of shared HCS with a cluster *c*_2_ (larger than *c*_1_) and the fraction of HCS shared is higher than *T*_3_, then *c*_1_ is merged into *c*_2_ and the HCS of *c*_1_ are added to the larger cluster. After a full iteration over all clusters, the cluster merging algorithm starts a new iteration, since some clusters and their associated HCS have now been updated. The algorithm terminates when no clusters are merged during an iteration. In a given iteration, if a cluster *c*_1_ can be merged into another cluster *c*_2_ which, in turn, can be merged into a third cluster *c*_3_, we first perform the merge operation of adding *c*_2_ to *c*_3_, postponing the merge of *c*_1_ into *c*_3_ (now also containing the HCS from *c*_2_) to the next iteration.

The intent with the cluster merging step is to merge clusters that stem from the same gene family but were divided into several clusters during the main clustering step due to not sharing enough HCS. This is mainly necessary for noisy data where there are not enough shared HCS due to, e.g., errors or sequencing artifacts. Additionally, HCS from new information in a cluster (SNPs or exons) could make a read pass the threshold *T*_2_ to be included in the cluster after it was visited in the initial clustering.

### 2.3 Input, output, and implementation details

The input of our algorithm consists of long transcriptomic reads that have been generated by either PacBio or ONT sequencing and were pre-processed via a barcode removal tool, such as Pychopper version 2.5.0 (for ONT data) and Lima (for PacBio data) removing artificial concatamers and non-full-length reads. Similarly to isONclust, isONclust3 outputs a tab separated file, which indicates the mapping of each read to a cluster, as well as separate *fastq* files containing the reads per cluster.

In our benchmarks we set *k* = 15 and *w* = 51 for PacBio and *k* = 13 and *w* = 21 for ONT, which is similar to isONclust (*k* = 15, *w* = 50 for PacBio and *k* = 13, *w* = 20 for ONT). We had to increase the value of *w* by 1 due to even values of *w* not being allowed in the minimizer implementation we use. The isONclust3 algorithm is written in the Rust programming language and is available via https://github.com/aljpetri/isONclust3. We use rust-bio Köster (2015) to parse the reads and the minimizer-iter library Martayan (????) for the generation of minimizers and canonical minimizers. We used rustc-hash, as the hash function for minimizers. IsONclust3 is not parallelized like other algorithms such as isONclust. A similar parallelization strategy to isONclust could be implemented, although the clustering is not guaranteed to be the same when using different amounts of cores.

## 3 Results

### 3.1 Deciding values for thresholds *T*_1_ and *T*_2_

We performed an experiment to assess suitable values of *T*_1_ and *T*_2_ for clustering (cluster merging deactivated) PacBio and ONT by analyzing one ONT (Droso) and one PacBio (ALZ) dataset. Following the results from the threshold analysis (Fig. 1 top row panels) we set *T*_1_ to 0.95 and 0.98 for ONT and PacBio data, respectively. Following the *T*_2_ threshold experiment (Fig. 1, bottom row panels), we set *T*_2_ to 0.5 for both ONT and PacBio data. For the cluster merging parameter *T*_3_, we also looked at the *V, c, h*, and *ARI* metrics for the Droso and ALZ datasets and set *T*_3_ to 0.5 and 0.8 for ONT and PacBio, respectively. One of the reasons that PacBio require a much higher *T*_3_ threshold is the presence of artificial concatamers (multiple transcripts present in the same read), which is a sequencing artifact that PacBio’s software, Lima, typically removes. We noticed many such artificial concatamers in the ALZ dataset, a dataset produced when Lima did not have the functionality to remove artificial concatamers implemented. Such artificial concatamers can yield over-clustering.

**Fig. 1:**
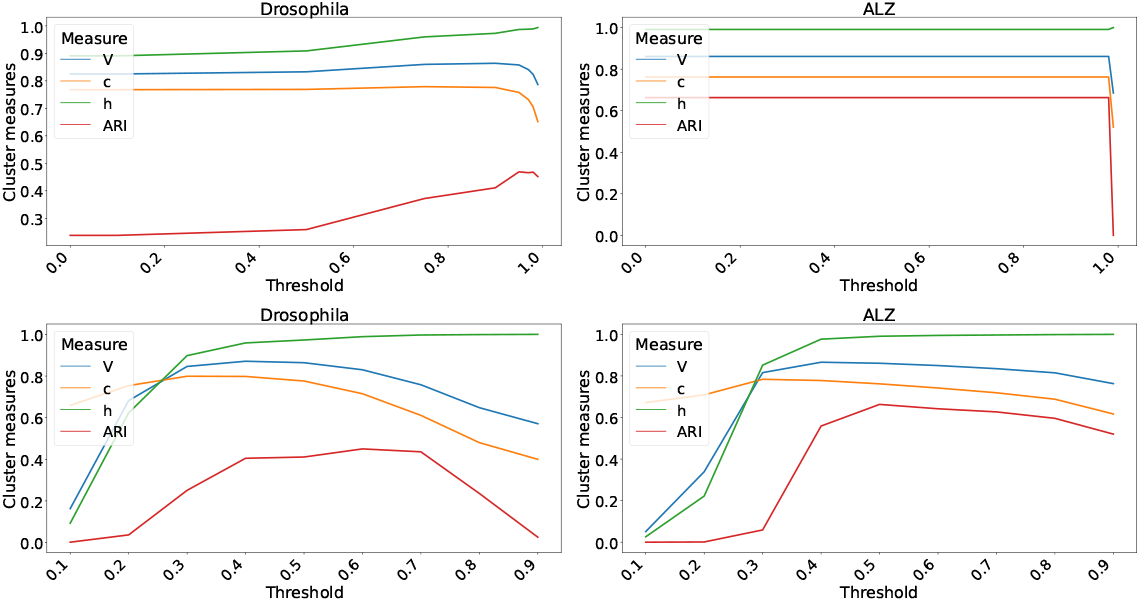
Threshold analysis for *T*_1_ (top row) and *T*_2_ (bottom row) on the ONT Droso (left panels) and a Pacbio ALZ (right panels) datasets. The x-axis indicates the value of *T*_1_ or *T*_2_, while the y-axis shows the resulting *V, c, h* and ARI values. For *T*_1_, we searched in [0, 0, 0.1, 0.5, 0.75, 0.9, 0.95, 0.97, 0.98, 0.99]. For *T*_2_ we searched in [0, 1, 0.2, …, 0.9].

### 3.2 Tools and datasets included in benchmark

We benchmarked isONclust3 against the existing state-of-the-art whole transcriptome clustering methods, namely isONclust, isONclust2, GeLuster, and RATTLE, on biological, synthetic, and simulated datasets (Table 1). We benchmarked the algorithms on three PacBio Human datasets that we call ALZ, PB_human_SIRV, and HG002 (details in suppl. Section S3), three mid-to-low error-rate ONT datasets from Drosophila (Droso), human (ONT_human), and a SIRV dataset with reads generated from a pool of known transcript products. We also included a simulated dataset and a high error-rate ONT dataset (ONT_old) used in the isONclust study Sahlin and Medvedev (2020). See Suppl. Section S3 for data availability.

**Table 1.** Overview over the datasets used for the evaluation: NS: non-singleton clusters (clusters containing more than one read), S: singleton clusters (clusters containing a single read), *: for the ONT_human dataset we estimated the average error rate using the Phred score *Q* = 13.5 (Ni et al. (2023)). We then calculated the respective error rate according to the Phred formula 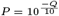.

| Dataset | Seq. Tech. | Reads |  |  | Classes (Estimated ground truth) |  |  |
| --- | --- | --- | --- | --- | --- | --- | --- |
|  |  | Error rate (%) | #reads | Unaligned (%) | NS | S | % reads in NS classes |
| SIRV | ONT | 6.9 | 1,409,565 | 1.1 | 7 | 0 | 98.8 |
| Droso | ONT | 7.0 | 3,764,978 | 13.0 | 5,488 | 2,153 | 86.9 |
| ONT_human | ONT | 4.5 * | 1,473,632 | 27.3 | 13,831 | 15,300 | 71.6 |
| ONT_old | ONT | 12.9 | 890,503 | 4.27 | 14,863 | 13,665 | 94.2 |
| SIM | Simulated | 1.9 | 500,000 | 4.0 | 14,792 | 2,152 | 99.6 |
| ALZ | PacBio | < 1.0 | 4,277,293 | 0.1 | 17,947 | 12,573 | 99.5 |
| PB_human_SIRV | PacBio | < 1.0 | 5,000,000 | 0.0 | 13,401 | 11,658 | 99.8 |
| HG002 | PacBio | < 1.0 | 37,200,651 | 0.0 | 27,055 | 45,198 | 99.9 |

### 3.3 Evaluation metrics

For the clustering quality assessment we use reference-guided clustering based on alignments to the reference genome via minimap2 Li (2018) as a proxy, similarly to previous studies Sahlin and Medvedev (2020); Ma *et al*. (2024). The *class* (ground truth estimate) of a read is the group to which the read was assigned by alignment. We denote reads as *unclassified* if the read could not be aligned. While the usage of our proxy of the true clustering may contain misclassifications through misalignments or sequencing artifacts in reads, e.g., due to reverse transcription errors, it is generally agreed upon as benchmarking approach for such methods and was used in previous studies Sahlin and Medvedev (2020); Ma *et al*. (2024). We consider it a reasonable way to assess the relative performance of clustering tools. Table 1 shows an overview of the alignment characteristics and classes of the datasets. We refer to clusters and classes as non-singleton (NS) if they contain more than one read and singleton (S) it they consist of only one read. The class classification serves as the truth when we compute the clustering accuracy of the algorithms.

As in Sahlin and Medvedev (2020); Ma *et al*. (2024), we used completeness *c*, homogeneity *h*, and *V* -measure Rosenberg and Hirschberg (2007), as well as the Adjusted Rand (*ARI*) statistics to assess the clustering quality of the algorithms. A cluster is homogeneous if all its clusters contain only data points from a single class. A clustering is complete if, for all classes, all the data points from a class are assigned to a single cluster. Intuitively, homogeneity penalizes wrongly clustering together reads from different classes, or over-clustering, while completeness penalizes reads that belong to the same class ending up in different clusters, or under-clustering. Specifically, let *X* be an array of *n* ordered reads where the elements in the array are the cluster identities (*c*_*id*_) as appointed by the clustering algorithm. We define *Y* in a similar manner for the ground truth class identities of the reads. The homogeneity measure is then calculated as *h* = 1 *− H*(*Y* |*X*)*/H*(*Y*). Here *H*(*∗*) and *H*(*∗*|*∗*) refer to the entropy and conditional entropy functions as described in Rosenberg and Hirschberg (2007). The completeness measure is given by *c* = 1 *− H*(*X*|*Y*)*/H*(*X*). The *V* -measure Rosenberg and Hirschberg (2007) is computed as the harmonic mean of *h* and *c*, and therefore combines the two measures in a way similar to the way precision and recall are combined into the *F* -measure for a classification problem. We also compute the Adjusted Rand Index (*ARI*) Hubert and Arabie (1985) to avoid bias with respect to a single clustering measure. Rand index (RI) as a measure of the percentage of correct pairings of elements. That is, a pairing (*a, b*) of elements *a* and *b* is correct if it agrees between the clustering and the classes, regardless of whether *a* and *b* belong to the same class or not, using the formula:

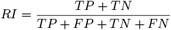

Where TP, TN, FP, and FN stand for True Positives, True Negatives, False Positives, and False Negatives, respectively. Depending on the nature of correct classes (e.g., the extremes of all singleton classes or all elements in one class), the expected similarity of a random clustering differs. The ARI is the RI but corrected for chance by subtracting the expected similarity of all pair-wise comparisons between clusterings specified by a random model. While the RI ranges *∈* [0, 1], the ARI can take values *∈* [*−*1, 1], where random labelings get an ARI close to 0. The *ARI* is calculated as follows:

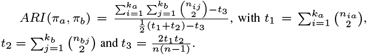

Here *π*_*a*_ and *π*_*b*_ are two clusterings of a dataset with *k*_*a*_ and *k*_*b*_ clusters, having *n* samples; *n*_*ij*_ indicates the number of common objects in cluster *c*_*i*_ in clustering *π*_*a*_ and in cluster *c*_*j*_ in clustering *π*_*b*_, *n*_*ia*_ denotes the number of objects in cluster *c*_*i*_ and *n*_*bj*_ represents the number of objects in cluster *c*_*j*_ in clustering *π*_*b*_. We also measured the percentage of reads in the data that was included in nontrivial clusters. As for most downstream applications (such as error correction or isoform prediction) minimal read numbers of 3-5 reads are necessary; non-clustered reads are discarded.

### 3.4 Benchmarking details

We ran isONclust3 with one core and all other algorithms with one and eight cores. For some algorithms such as isONclust and isONclust2, both clustering quality and memory usage will change depending on the number of cores. We ran isONclust3 both with and without canonical minimizers, denoted isONclust3 and isONclust3_nc) respectively, to illustrate the impact on datasets where a gene expresses transcripts from both directions. We also ran isONclust3 with cluster merging (with canonical minimizers) to indicate its effect on the final clustering, denoted isONclust3_cm. We evaluated wall clock time using `usr/bin/time/time -v`. To minimize the impact of file access times on the overall run times, we let all algorithms write their output onto fast access scratch memory space, a space available individually for each compute node. We used a CentOS 7, with two 10-core Intel Xeon V4 CPUs each. The analyses were run on the default nodes having 128GB of memory and high-RAM nodes, each having 256GB of RAM available, when needed.

### 3.5 Clustering quality evaluation

Overall, isONclust3 performs favorably or comparable to the other tools, especially when running with cluster merging enabled (Fig. 2). The only exception to this was the ONT_old dataset (avg. error rate 12.9%) where isONclust3 clusters fewer reads than some of the other algorithms, suggesting that different thresholds might be better for datasets with very a high error rate. While all algorithms have some dataset where they perform well, isONclust3 and GeLuster algorithms typically show better results than isONclust, isONclust2 and RATTLE. The *V, c* and *h* measures show similar results for isONclust3 and GeLuster, but isONclust3’s *ARI* is consistently higher for all the datasets. Also, for the largest dataset, HG002, which was produced with the new PacBio Revio system, none of the algorithms except isONclust3 was able to run on our server due to too high RAM usage (*>* 256Gb).

**Fig. 2:**
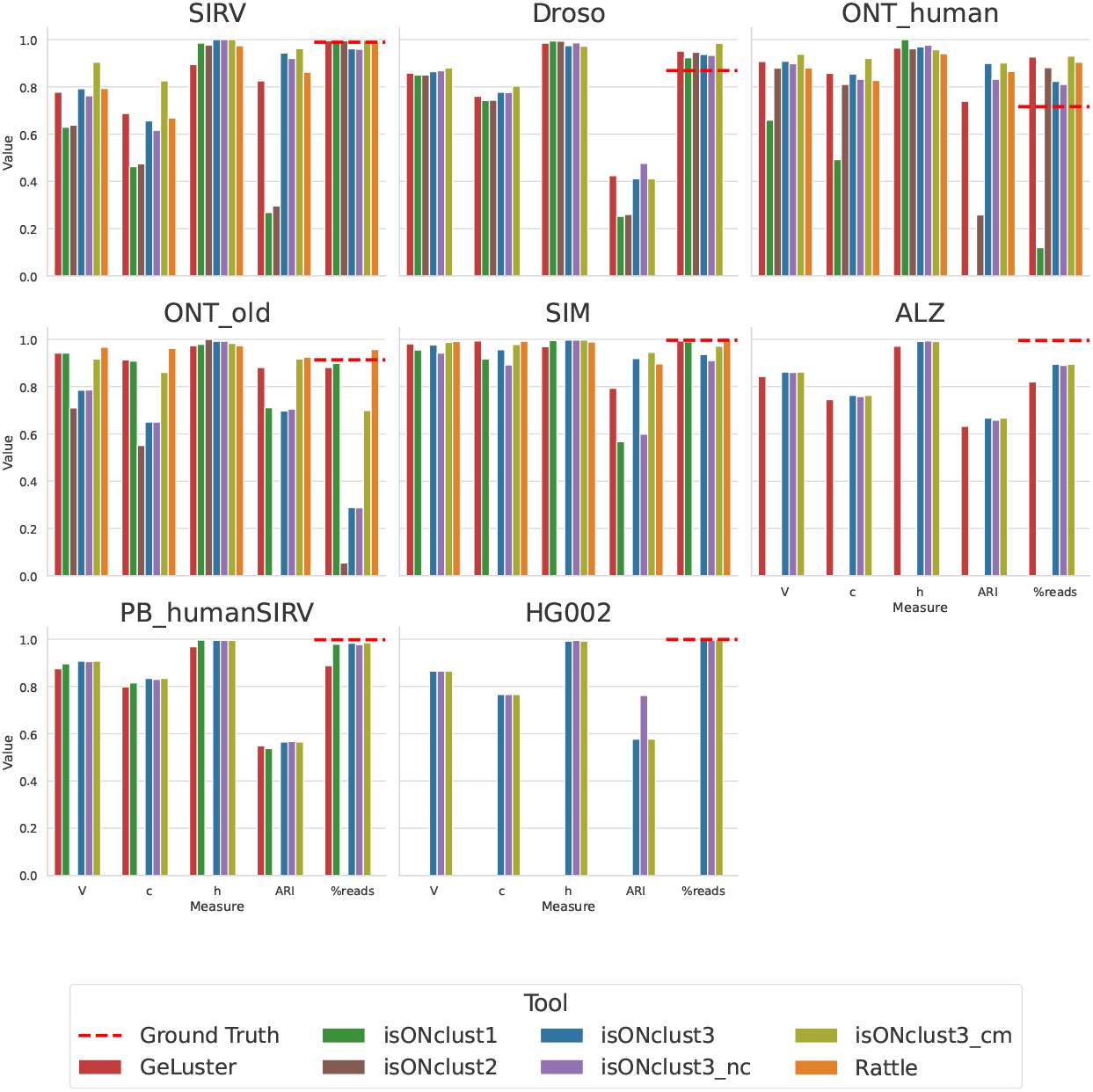
Cluster quality measures using one core. For the ALZ dataset isONclust and RATTLE did not finish during the given run time limit (120 hours). For the HG002 dataset no algorithm except isONclust3 was able to be run due to their RAM usages exceeding the cluster RAM usage limit of 256GB. The dashed red line (labeled ground truth) shows the fraction of reads that were placed into non-singleton classes, which is merely an indicator for the amount of reads that can be clustered, subjects to the limitations of the alignment based ground truth estimation as mentioned in section 3.2.

As for the effect of cluster merging, isONclust3_cm improves *V, c*, and *ARI* over isONclust3 on the ONT datasets, but did not improve clustering for the PacBio datasets, indicating that cluster merging is beneficial primarily for the noisier datasets.

Using canonical or non-canonical seeds impacts clustering of reads expressed from the same genes but in different directions. IsONclust3_nc yields results close to the canonical version except for the SIRV and SIM datasets which contain genes expressed in both directions. Notably, isONclust3_nc shows better results than isONclust (which uses non-canonical seeds). Together with the fact that we use similar parameter values of *k* and *w* in isONclust3, it indicates the improvements in isONclust3 over isONclust come from the algorithmic differences (usage of HCS and dynamic updating). For the ONT_human dataset, isONclust was only able to cluster 12 percent of the reads. We did not observe significant differences in clustering quality when the other tools were run with 8 cores instead of one core (Suppl. Fig. 2).

IsONclust3 and GeLuster are also more reliable when it comes to finishing the clustering out of the five benchmarked algorithms. IsONclust2 could only be run on the ONT datasets. On this data, isONclust2 showed worse results compared to isONclust3. RATTLE terminated with a segmentation fault on the Droso and did not finish within 120 hours on the ALZ dataset when using both one and eight cores. None of the algorithms except isONclust3 could run on the HG002 dataset, regardless of whether one or eight cores were specified.

### 3.6 Runtime analysis and memory usage assessment

Figure 3 shows runtime and peak memory usages of all algorithms when using one core. Missing datapoints indicate that the algorithm either ran out of memory (256Gb) or took more than the given time limit of five days (120h). IsONclust3 consistently has the lowest runtime and is 10 to 100 times faster than isONclust, RATTLE, and GeLuster on individual datasets. IsONclust2 is comparable to the runtime of isONclust3 on the ONT datasets, but its memory usage exceeded the memory usage of any other tool we benchmarked. We also ran the other algorithms with eight cores and included isONclust3 with one core as a comparison (Suppl. Fig. 2). When the other algorithms are run with eight cores, isONclust3 is still 5 to 20 times faster for larger, challenging datasets such as ALZ. Notably, isONclust3 is the only algorithm that can cluster the HG002 dataset on our system due to the high memory usages of other algorithms, regardless of the number of cores used. For the ALZ and PB_humanSIRV datasets RATTLE exceeded the time limit. IsONclust was able to complete clustering of ALZ in the time limit only when run with eight cores. The GeLuster algorithm showed a high runtime for the ALZ and PB_humanSIRV datasets, which we did not observe for the isONclust3 algorithm. In terms of memory usage, isONclust3 used similar or slightly more RAM than GeLuster on all datasets except on the largest dataset, HG002, where GeLuster failed. Of the datasets in which both tools completed, the largest discrepancy was on ALZ where GeLuster used 25.6Gb and isONclust3 used 37.0Gb.

**Fig. 3:**
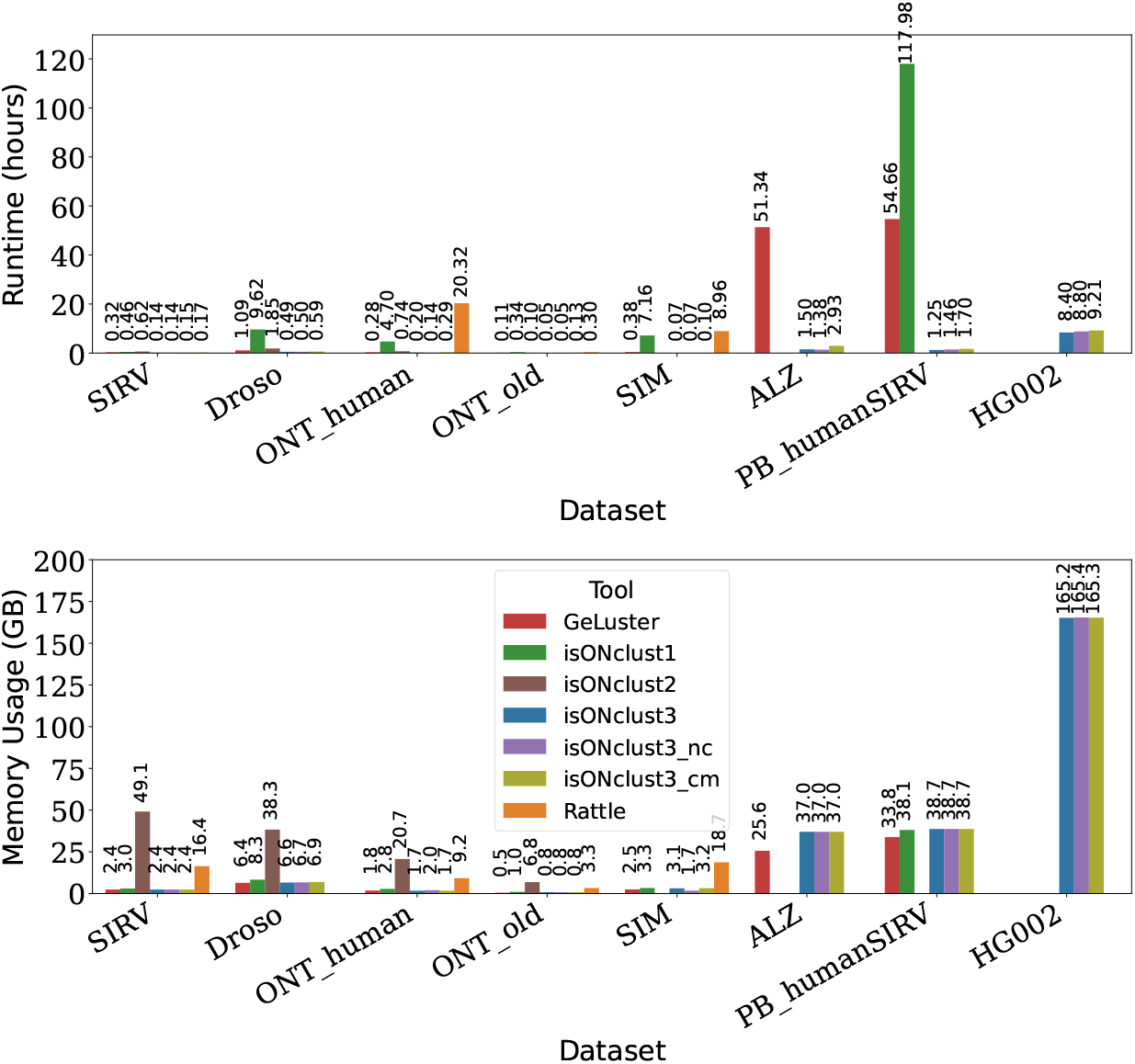
Runtime and memory usage. Upper panel: runtime of each algorithm for each dataset in hours. Lower panel: memory usage in GB. x-axis: the different datasets and tools, y-axis: Runtime in hours/ RAM usage in GB. IsONclust3 was the only algorithm able to run the HG002 dataset on our cluster.

IsONclust3_cm increases overall runtime of isONclust3. We observed the highest runtime increase for ONT_old (161%). This meant an increase of overall runtime from 3 minutes to almost 8 minutes. The peak RAM usage of the isONclust3 algorithm is not affected by the cluster merging step as the reads do not have to be stored during this step. IsONclust3 with one core also remains faster than other tools using eight cores (Suppl. Fig 3).

## 4 Discussion

We introduced the isONclust3 algorithm for clustering massive transcriptomic long-read sequencing datasets. IsONclust3 uses minimizer seeds to compare reads to clusters and dynamically updates representations of clusters with high-confidence minimizer seeds instead of choosing seeds from one read as the representative of a cluster (isONclust). Furthermore, isONclust3 adds an iterative cluster merging step to its algorithm to improve the clustering at low runtime overhead. The isONclust3 algorithm is also implemented in the high-performance programming language Rust also facilitating simple installation and usage of the tool (unlike the Python implementation of isONclust and the prototype isONclust2). We showed that isONclust3 performs favorable to existing tools in terms of cluster measures on a diverse set of datasets. For noisier datasets (e.g., the ONT datasets in our benchmarks), isONclust3’s cluster merging improves cluster quality further, while for PacBio datasets we did not see any significant improvement. This is likely because with noisier data, it takes more time to find HCS to represent the cluster which motivates another pass with a more complete set of HCS. We therefore recommend the usage of the cluster merging for ONT data.

IsONclust3 uses canonical minimizers as default seeds. However, to enable a better head-to-head comparison to isONclust, we also ran isONclust3 with noncanonical minimizers, showing the algorithmic improvements, beyond canonical seeds, over isONclust. IsONclust3 also outperforms all other tools in memory usage and runtime on most datasets, and is the only tool that can cluster PacBio Revio sequencing datasets with ten times more reads to older sequencing machines. IsONclust3 replaces isONclust, and is applicable to any use case that isONclust has been utilized for.

### 4.1 Future work

While isONclust3 generally outperforms existing algorithms with respect to runtime and RAM usage, it is not yet parallelized. If parallelized in the same manner as isONclust, it could improve the runtime of isONclust3 further with a trade-off in RAM usage (depending on how many cores are used). There are two memory bottlenecks in isONclust3 due to all reads being present in RAM at the same time. The first is in the sorting step of the reads, and the second is in the output step when writing reads to clusters. It is possible to use, e.g., external merge sort to sort the reads to alleviate the first bottleneck. The second could be resolved with a similar external memory algorithm. However, such algorithms will result in a time-to-memory trade-off, and we consider it future work to optimize this. We may also consider using bit-encoding of two bits per nucleotide to represent the reads, and possibly discretizing the Phred quality values further to fit in less bits. As for the methodology aspects, strobemers Sahlin (2021) have been used successfully in de novo transcriptome analysis tools to classify transcripts Nip *et al*. (2023) based on a set of overlapping strobemers. It is possible that isONclust3 could implement such an approach to compare and cluster reads on transcript-level for applications that require it, or to further increase processing speed through uniqueness of the seeds.

## 5 Conclusion

We presented isONclust3, an algorithm for clustering long-read transcriptomic data by gene families. Using simulated, synthetic, and biological data we demonstrated that isONclust3 produces accurate clusterings and has comparable or higher *V, c, h* and *ARI* than other methods. IsONclust3 is also substantially faster and less memory consuming than other tools on most datasets and is the only algorithm that can cluster a PacBio Revio dataset with 37M reads. IsONclust3 replaces isONclust, and is suitable to cluster recent transcriptome long-read sequencing datasets with tens of millions of reads.

## Availability

IsONclust3 is available at https://github.com/aljpetri/isONclust3. The analysis pipelines were written using snakemake Mölder *et al*. (2021). The pipeline and intermediate results used for plotting are available via https://github.com/aljpetri/isONclust_analysis. The sources of the datasets can be found in the supplementary materials to this paper.

## Acknowledgements

Parts of the evaluation were enabled by resources in project snic2022-5-592 provided by the Swedish National Infrastructure for Computing (SNIC) at UPPMAX, partially funded by the Swedish Research Council through grant agreement no. 2018-05973, and by resources at the PDC Center for High Performance Computing, KTH Royal Institute of Technology, partially funded by the Swedish Research Council through grant agreement no. 2016-07213.

