## Supplementary file for "De novo clustering of large long-read transcriptome datasets with isONclust3"

#### Contents

**S.1Pseudocode of the sorting and clustering steps** **SI-2**

**S.2Time complexity analysis of isONclust3** **SI-2**

**S.3Data availability** **SI-3**

**S.4Evaluating the effect of different values of $k$ and $w$ on the clustering** **SI-3**

**S.5Additional figures and tables** **SI-4**

#### S.1 Pseudocode of the sorting and clustering steps

```
input  : Reads R, k, w, thresholds T1, T2
output: Read clusters C

Function SortReads(R, k, w, T1)
    for r ∈ R do
        seeds = generateSeeds(r.sequence, k, w)
        hcs = findHCS(seeds, T1)
    end
    R_sort ← R.sort(hcs)
    return R_sort

Function Clustering(R_sort, T1, T2)
    Initialize hash table C (clusterID → readIDs).
    Initialize hash table S (seed → clusterIDs).
    c_id = 0
    for r ∈ R_sort do
        seeds = generateSeeds(r.sequence, k, w)
        if C is empty then
            C.insert(c_id, r.acc)
            hcs = findHCS(seeds, T1)
            S.addNewHCS(hcs)
            c_id += 1
        end
        else
            Initialize hash table H (clusterID → seed count)
            for s ∈ seeds do
                H ← SeedCount(s, S, T2)
            end
            for c ∈ clusters do
                bestCluster ← compareReadToCluster(r, c.representative, T2)
            end
            A, c' = max(H.counts, H.clusterID )
            hcs = findHCS(seeds, T1)
            if A/|r.seeds| > T2 then
                C[c'].add(r.acc)
                S.addNewHCS(hcs, c')
            end
            else
                C.insert(c_id, r.acc)
                S.addNewHCS(hcs, c_id)
                c_id += 1
            end
        end
    end
    return C
R_sort = SortReads(R, k, w, T1)
C = Clustering(R_sort, T1, T2)
```

**Algorithm 1:** isONclust3 sorting and clustering.

#### S.2 Time complexity analysis of isONclust3

Our algorithm is a greedy heuristic and therefore it is challenging to derive an informative worst-case runtime. We attempt to give a worst case analysis by parameterizing our analysis. Treating the function for hashing k-mers $h$ as a constant and letting $l$ be the length of the longest read, then producing minimizers has time complexity $O(hln)$ for $n$ reads while filtering of minimizers to yield HCS takes at most $O(kln)$. The sorting step is $O(n \log n)$ for $n$ reads (we use timsort).

As for the clustering step, the worst case scenario is that all reads form new clusters and the fraction of shared seeds with the HCS of existing clusters is high but lower than $T_2$. If we let $T_2 > 1$ (forcing no clustering) with a set of identical reads where all seeds $(< l)$ are shared, the clustering has time complexity $O(ncl)$, since each of the $n$ reads need at most $l$ seed look-ups and updates at most $c$ cluster identifier counters for each seed. The runtime is therefore $O(hln + kln + n \log n + ncl))$. In the worst case of $n = c$ (each read forms a new cluster), this is dominated by $O(ncl)) = O(n^2 l)$. However, in practice there are substantially less clusters than reads, and seeds typically only match a small subset of clusters.

The worst case scenario for the iterative cluster-merging step would be that precisely one cluster is merged into another cluster in each iteration. Each cluster-merging iteration has the same complexity as the clustering step. Since there is at most $c$ iterations for the cluster merging in this scenario, the cluster merging is in worst case $O(n(hln + kln + n \log n + c^2 l)))$. However, it is in theory impossible to have the two cases that no reads are initially clustered and the post-merging is merging clusters true at the same time. So the upper bound is not tight.

#### S.3 Data availability

We downloaded the ONT_human dataset from the NCBI database with accession DRX524696 (`https://www.ncbi.nlm.nih.gov/sra/DRX524696`). The SIRV and Drosophila data are available via the ENA browser with the project accession number PRJEB34849. Drosophila reference genome (assembly BDGP6.22) was downloaded from `ftp://ftp.ensembl.org/pub/release-97/fasta/drosophila_melanogaster/dna/Drosophila_melanogaster.BDGP6.22.dna.toplevel.fa.gz`. We used the Ensembl release 97 annotated on assembly BDGP6.22 for the Drosophila data, obtained from `ftp://ftp.ensembl.org/pub/release-97/gtf/drosophila_melanogaster/Drosophila_melanogaster.BDGP6.22.97.gtf.gz`. The SIRV genes and gene annotations were downloaded from `https://www.lexogen.com/wp-content/uploads/2018/08/SIRV_Set1_Lot00141_Sequences_170612a-ZIP.zip`. We used the T2T-CHM13v2.0 assembly that we downloaded from `https://github.com/marbl/CHM13` as human reference for the ALZ and ONT_human experiments. We downloaded the ALZ dataset from `https://downloads.pacbcloud.com/public/dataset/Alzheimer2019_IsoSeq/`. We downloaded the PB_human_SIRV dataset from `https://downloads.pacbcloud.com/public/dataset/UHRRisoseq2021/Intermediate-FullLengthReads/`. We used the first 5 million reads of this dataset to enable all tools to run on it. The dataset is sequencing read from a human reference with SIRV spike-ins added. To generate the reference for the data we concatenated the human reference with the SIRV reference files. The HG002 dataset was downloaded from `https://downloads.pacbcloud.com/public/dataset/Kinnex-full-length-RNA/DATA-Revio-HG002-1/2-FLNC/`. We obtained the high error rate ONT dataset ONT_old from `https://s3.amazonaws.com/nanopore-human-wgs/rna/fastq/Bham_Run1_20171115_1D.pass.dedup.fastq`.

#### S.4 Evaluating the effect of different values of $k$ and $w$ on the clustering

In the main paper analysis we used the same parameters of $k$ and $w$ as isONclust ($k = 13$, $w = 20$, except$w$ was increased by 1 due to limits of a third party library). This was

| k | w | V | c | h | ARI |
|---|---|---|---|---|---|
| 11 | 17 | 0.905 | 0.839 | 0.981 | 0.829 |
| 11 | 19 | 0.902 | 0.834 | 0.982 | 0.828 |
| 11 | 21 | 0.605 | 0.792 | 0.489 | 0.025 |
| 11 | 23 | **0.909** | **0.848** | 0.98 | 0.832 |
| 11 | 25 | 0.906 | 0.842 | 0.981 | 0.829 |
| 13 | 17 | 0.883 | 0.801 | 0.985 | 0.810 |
| 13 | 19 | 0.881 | 0.797 | 0.985 | 0.807 |
| 13 | 21 | **0.908** | **0.854** | 0.969 | **0.899** |
| 13 | 23 | 0.877 | 0.79 | 0.985 | 0.798 |
| 13 | 25 | 0.874 | 0.785 | 0.985 | 0.793 |
| 15 | 17 | 0.86 | 0.763 | 0.987 | 0.769 |
| 15 | 19 | 0.857 | 0.757 | 0.987 | 0.755 |
| 15 | 21 | 0.892 | 0.822 | 0.975 | **0.903** |
| 15 | 23 | 0.851 | 0.748 | **0.988** | 0.745 |
| 15 | 25 | 0.848 | 0.742 | **0.988** | 0.737 |

**Table 1.** Overview for different values of $k$ and $w$. Top two results in each column are bold faced. The results in this table indicate $k = 13$, $w = 21$ to have the best overall combination of $V$ ,$c$, $h$ and ARI.

done to investigate relative performance of the algorithms for the same parameter setting. However, we also tested the effect of other parameter choices on the ONT_human dataset. For this we ran isONclust3 with any combination of $k = \{11, 13, 15\}$ and $w = \{17, 19, 21, 23, 25\}$ and evaluated the results with respect to the cluster quality measures $V$, $c$, $h$ and ARI. The results of this analysis can be found in table 1.

### S.5 Additional figures and tables

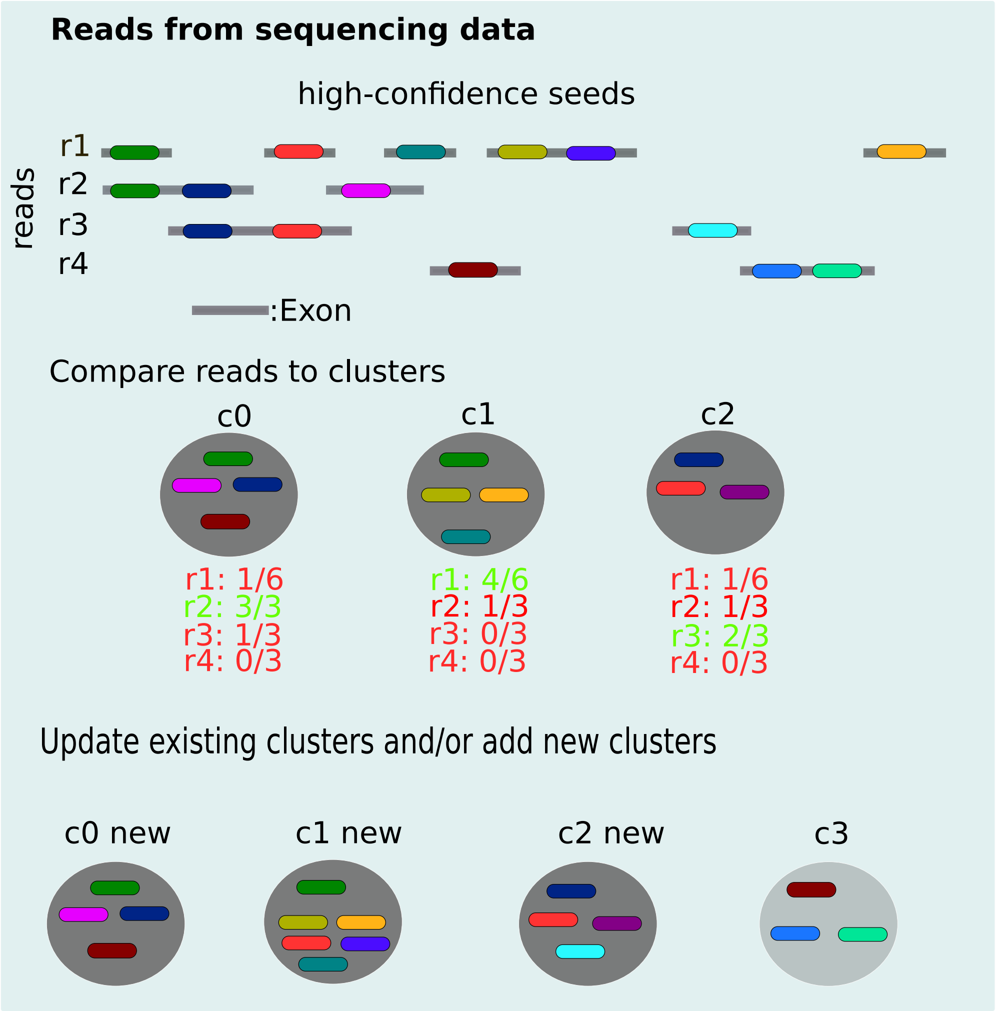


**Figure 1. Visualization of the dynamic clustering approach:** Top: A set of reads with seeds (colored), where in this example, all seeds are also HCS. Each line represents an exon, which may span over several seeds. Middle: a set of existing clusters that the reads are compared to. If the portion of HCS in the read shared with a cluster $c_i$ exceeds $T_2$ (0.5), the read is appointed to the cluster. Lower panel: Updating of the clusters: In our example, read r2 is fully contained in cluster c0. It is therefore added to the cluster without having to update the cluster representative. Read r1 has a high enough overlap to be added to cluster c1, therefore the cluster representative is updated with the HCS not previously present in it but in the read. The same is happening for read r3 and cluster c2. As read r4 does not have enough overlap with any of the existing clusters a new cluster, c3, is formed from it and its HCS.

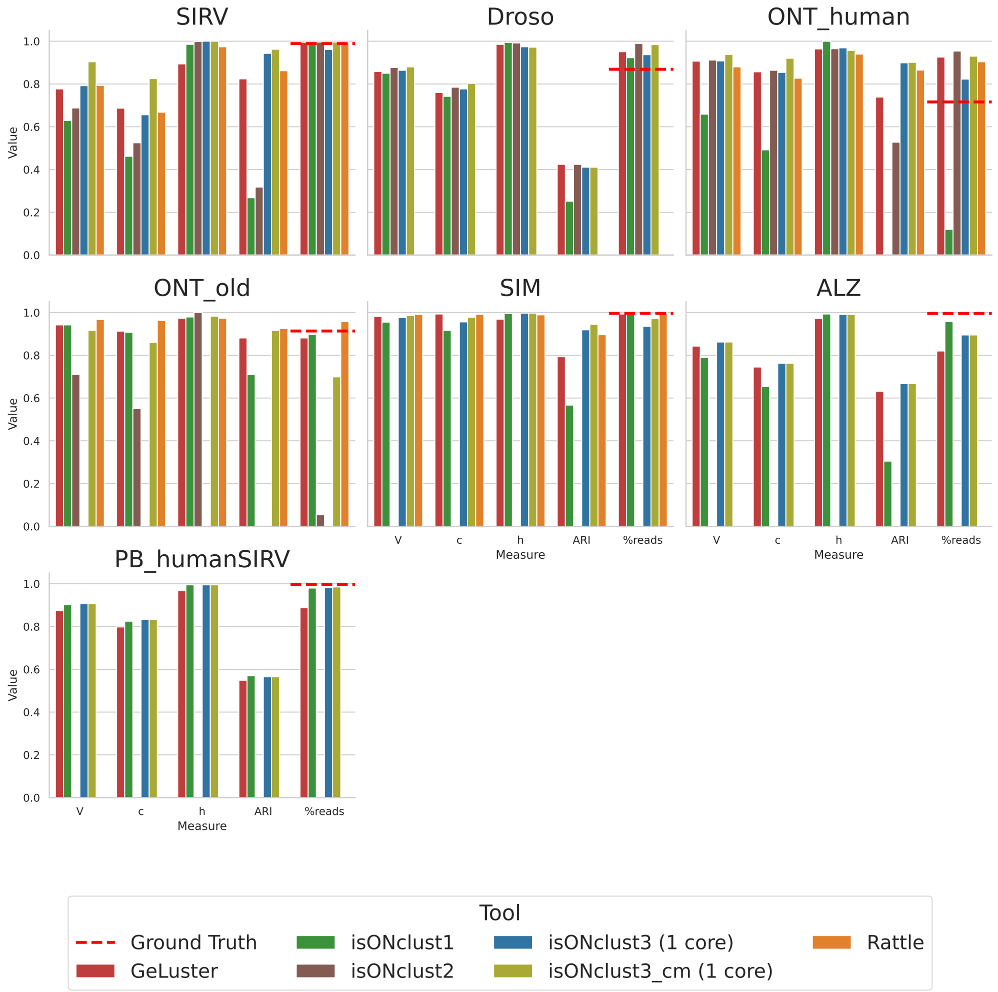


**Figure 2. Performance of the analyzed algorithms on the datasets run with eight cores:** Clustering measures for each dataset and each tool (x-axis: the cluster measure, y-axis: the respective resulting value)

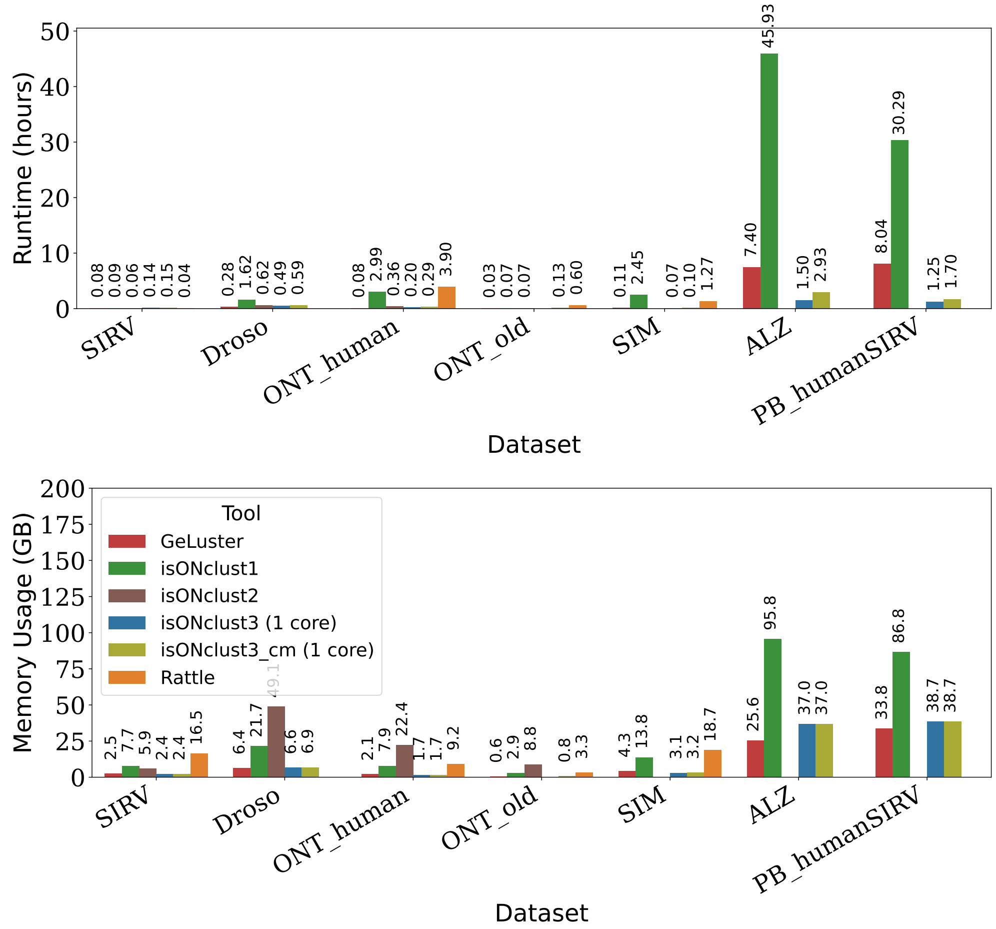


**Figure 3. Performance of the analyzed algorithms on the datasets run with eight cores:** Top panel: runtime of each algorithm for each dataset in hours, lower panel: memory usage in GB. x-axis: the different datasets and tools, y-axis: Runtime in hours/ RAM usage in GB. Please note that the isONclust3 algorithm was run with 1 core but added to this figure to enable a better visual comparison.
